# Thermoplasmonic induced vesicle fusion for investigating membrane protein phase affinity

**DOI:** 10.1101/2022.09.19.508467

**Authors:** Guillermo Moreno-Pescador, Mohammad R. Arastoo, Salvatore Chiantia, Robert Daniels, Poul Martin Bendix

## Abstract

Many cellular processes involve the lateral organization of integral and peripheral membrane proteins into nanoscale domains. Despite the biological significance the mechanisms that facilitate membrane protein clustering into nanoscale lipid domains remain enigmatic. In cells the analysis of membrane protein phase affinity is complicated by the size and temporal nature of ordered and disordered lipid domains. To overcome these limitations we developed a method for delivering membrane proteins from transfected cells into phase-separated model membranes that combines optical trapping with thermoplasmonic-mediated membrane fusion and confocal imaging. Using this approach we observed clear phase partitioning following the transfer of GFP-tagged influenza hemagglutinin and neuraminidase from transfected cell membranes to giant unilamellar vesicles. The generic platform presented here allows investigation of the phase affinity of any plasma membrane protein which can be labeled or tagged with a fluorescent marker.

## Introduction

Lipid domains play important roles in organization of the cellular plasma membrane and hence control a number of processes (1, 2) ranging from membrane trafficking (3) to apoptosis (4, 5). They are also involved in a number of diseases including cancer emergence and invasion (6, 7), and cardiovascular diseases (8). Moreover, the discovery of a high level of cholesterol and saturated lipids in the envelope of HIV (9) together with the finding that ordered membrane domains play a role in pathogenic microorganisms (10) supports the hypothesis of a modulatory role of lipid domains in host-pathogen interactions (11) including virus entry (12) and budding (13).

The proliferation of many enveloped viruses is intimately dependent on the structure and organization of the viral proteins in the plasma membrane. For influenza viruses, clustering of hemagglutinin (HA) and neuraminidase (NA) in the plasma membrane is crucial for the assembly and budding of progeny virions. The mechanism behind the lateral organization of proteins in the plasma membrane remains enigmatic, but the plasma membrane lipids have been proposed to be responsible for recruitment of transmembrane proteins to nanoscale budding-sites of virus infected cells (14, 15). The idea of lipids being responsible for clustering virus proteins at the cell surface is appealing and indeed co-localization of viral envelope proteins, with massive ectodomain heads, could produce an asymmetric lateral pressure across the membrane thus causing bending of the membrane and contribute to the viral budding (16). Eukaryotic cell membranes do form dynamic lipid raft structures enriched with proteins and this allows cells to perform lateral organization of proteins into nanoscale domains (17). Such organization could be critical for cell functions considering that membrane proteins constitute up to one third of a mammalian cell proteome (18, 19) and that the plasma membrane contains roughly 30.000 proteins per *μ*m^2^ populating 30-55% of the membrane area (20). Enriched in cholesterol and sphingolipids, lipid-raft domains could provide sorting platforms for both transmembrane and peripheral proteins possibly including viral proteins (4, 21–27).

The highly dynamic structure (milliseconds) and very small size (10-200nm) of rafts makes visual detection and investigation of rafts difficult (28). The current methods for studying rafts are based on biochemical, biophysical, computational and analytical tools (29). Biochemical tools use detergents to solubilize membrane lipids and proteins (30) and do not properly reflect the native molecular structure and organization of rafts. Biophysical tools are mainly based on model membranes (31) including Giant Unilamellar Vesicles (GUVs) into which transmembrane proteins can be reconstituted using biochemical protocols (32, 33). Although computational tools provide insights into the molecular behavior with unprecedented precision (34), they are model driven and need to be experimentally validated. Analytical tools recruit a wide range of techniques from microscopy to label-free methods (29). Despite the high resolution provided by some assays, most analytical tools require sample preparation involving tedious procedures (35) which can potentially influence the protein localization in model membranes.

Transmembrane proteins from influenza virus have been studied using cell-derived Giant Plasma Membrane Vesicles (GPMVs) which can exhibit large scale phase segregation only at very low temperatures (~ 5°C) and by addition of the cross-linking agent choleratoxin B (32, 36). These studies have shown that neuraminidase and hemagglutinin do partition into the more disordered lipid phase which is somewhat conflicting with the idea that viruses bud from cholesterol enriched domains. These findings have opened up new questions on how virus budding and plasma membrane structure are related, thus raising the possibility that raft-association of virus proteins could be mediated through interactions between different proteins. However the low temperature, and use of a toxin to trigger phase separation, raises concerns whether this system sufficiently reflects the cellular plasma membrane.

Here, we present a general assay which can address the localization of transmembrane proteins in phase separated membranes at any relevant temperature without any chemical cross-linking agent. Phase separated GUVs are fused with GPMVs containing the protein of interest in the correct orientation. The fused hybrid vesicle contains phase separated domains which allow us to study the localization of membrane protein phase affinity at physiologically relevant temperatures. We demonstrate the simplicity and efficacy of the method by investigating the phase affinity of influenza virus A transmembrane proteins and several mutated versions of these proteins. Fusion of the vesicles is accomplished using thermoplasmonics and optical trapping (37–39). This generic method can be applied to any transmembrane protein expressed in cells and does not require protein purification. Finally, it also holds the potential for studying the interactions between either transmembrane proteins or between transmembrane and peripheral membrane binding proteins in a controlled manner under physiological conditions.

## Experimental Section

### In-vitro transmembrane protein delivery to the model membranes

Eukaryotic cells contain a variety of membrane proteins exhibiting diverse functions such as signaling, which may be dependent on raft association of proteins. The plasma membrane appears uniform at scales larger than the optical diffraction limit which means that any possible lipid domains must be smaller than ~ 200 nm making them difficult to study using confocal microscopy. To reflect the behavior of these proteins, we utilized thermoplasmonic fusion mediated by optically trapped gold nano-heaters to deliver raft-associated proteins into model membranes having clearly visible and ordered lipid domains of micrometer scale.

At first, the chimeric fluorescent protein of interest is transiently expressed in HEK293T cells to have the proteins in the physiological condition. Following expression a vesiculating reagent is added to the cells which triggers cells to detach part of their plasma membrane, containing the associated proteins, as Giant Plasma Membrane Vesicles (GPMVs) also known as blebs. The produced GPMVs hold the protein of interest with the correct orientation with intact functionality (36). To transfer the transmembrane proteins to a model membrane we mix the GPMVs with phase separated GUVs in a cell culture dish (Fig. 1A) and Fig. S1. The GUVs vary in size with a diameter between 10 - 80 *μ*m and are composed of saturated fatty acids plus cholesterol and unsaturated lipids. The stochiometry of lipids and cholesterol allows formation of stable liquid-ordered phase (L_o_, raft resembled domain) and the liquid-disordered phase (L_d_, non-raft resembled domain), respectively at room temperature (T = 22°C).

**Fig. 1.**
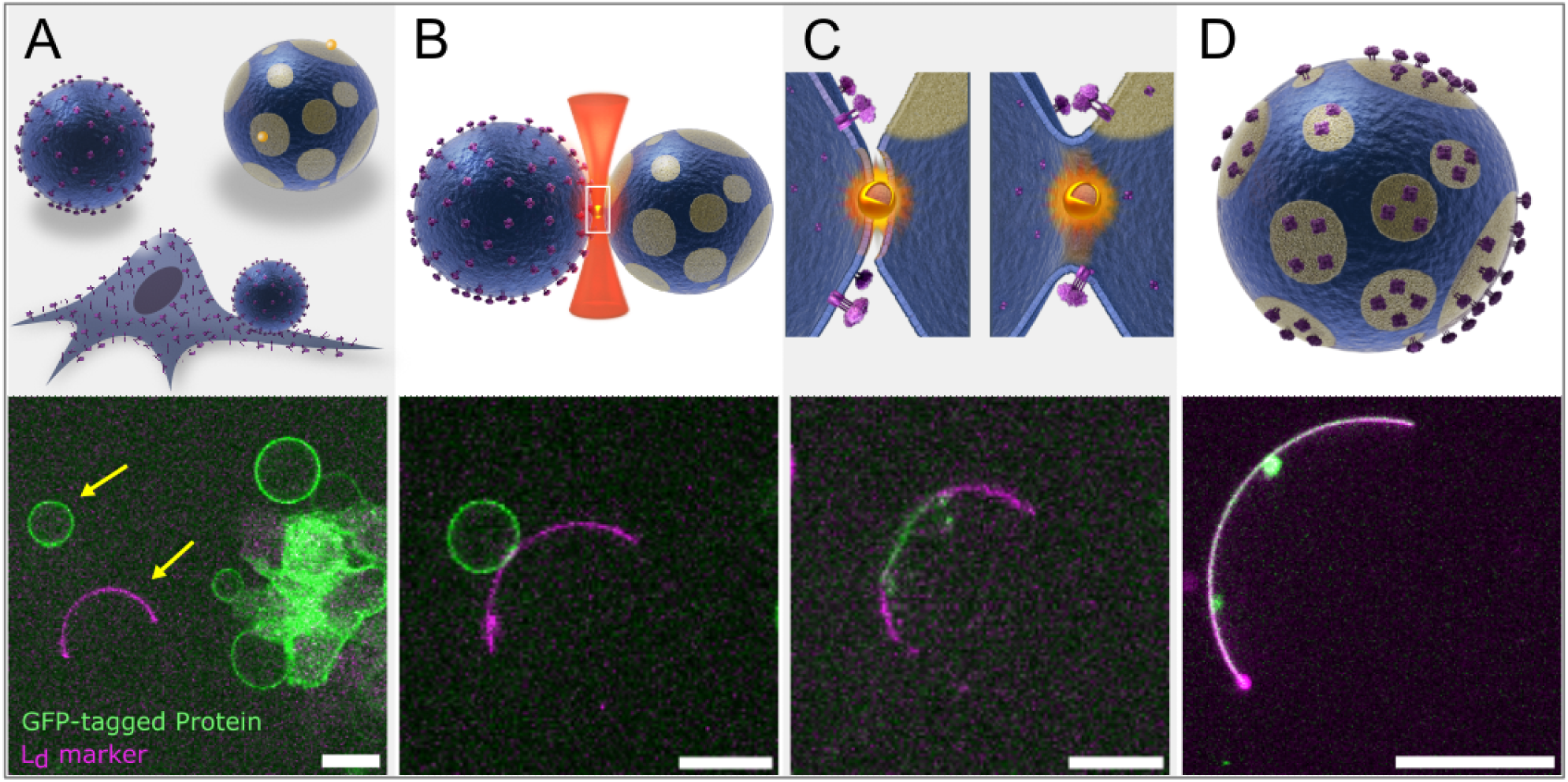
In vitro thermoplasmonics-based delivery of integral membrane proteins to phase-segregated model membranes. **A.** HEK293T cells are transfected to express the protein of interest, here GFP-tagged hemagglutinin Transmembrane Domain (HA-TMD) with the ectodomain labeled with a GFP. Further, they are treated with the blebbing reagent that results in the production of GPMVs containing the expressed transmembrane protein in its correct orientation. Phase separated Giant Unilamellar Vesicles (GUVs) are also added to the cell culture. As the fusion mediator, 150nm streptavidin conjugated Gold NanoShells (AuNSs) are linked to the GUVs through biotin-steptavidin interaction. Prior to fusion a GPMV is grabbed by the optical tweezers and brought to close proximity of a GUV (yellow arrows). **B.** Irradiation of the GPMV-GUV interface by the NIR laser brings a AuNS into the laser focus where it produces a local heat sufficient to fuse the apposing membranes. **C.** The process of lipid mixing following fusion. **D.** The resultant vesicle retains its ordered and disordered phases and the HA-TMD partitions into the liquid disordered phase (marked by 18:1 Liss Rhod. PE). Scale bars are 15 *μ*m.

Because the GPMVs also differ in size, variations in the GUVs diameter enables us to select GUV-GPMVs of comparable size for successful fusion (Fig. 1B). The NIR laser (*λ* = 1064nm) is then used to grab and position the GPMV of choice in close proximity to the selected GUV. To make the fusion non-random and more efficient, 150 nm streptavidin conjugated gold nano shells (AuNSs) are bound to biotinylated lipids in the GUV membrane. When the close contact between the GPMV and the GUV is established, the focus of the optical trap is positioned at the contact point of the two apposing membranes. The gold nanoparticles are mobile on the fluid GUV membrane and are pulled into the optical focus (Fig. 1C). Irradiation of the AuNS by the NIR laser, operated at 3.04 × 10^10^ W m^-2^ at the sample, produces highly localized heat (in the order of Δ*T* ~ 200K) (39) which is sufficient to transiently open the two apposing membranes and thereby, fuse the vesicles together to form a hybrid vesicle (Fig. 1D). AuNSs produce a thermal pulse when entering the optical focus and subsequently escape the focus or become structurally degraded (40) which significantly decreases their ability to absorb light. The transient (< 1s) and local (~ 100 nm) heating has the advantage of preventing thermal damage to lipids and the membrane associated proteins. The hybrid vesicle has the lipid composition originating from both the plasma membrane and the GUV lipid species. By this way, membrane associated proteins are also delivered to the phase separated model membrane where they can move freely and occupy the phase of their preference (Fig. 1E). Protein delivery to phase segregated model membranes by this method is not limited to membrane proteins, but it can also be extended to any relevant system that could be introduced in either of the components of this experimental assay.

We tested the assay on five different proteins and a control GPI anchor known to favor ordered membrane phases. Plasmids encoding for full length hemagglutinin A (HA-FL) and its transmembrane domains (HA-TMD) were engineered. Additionally, plasmids encoding for full length neuraminidase (FL-NA) and two mutants: the transmembrane domain NA-TMD (NA-42) encodes for the first 42 amino acids which primarily contains the transmembrane and short N-terminal domain and (NA-Δhead (NA-62) encodes for the first 62 amino acids, removing the large head domain and a portion of the stalk.

## Results and discussion

To demonstrate the applicability of this method we examined the influenza membrane proteins hemagglutinin (HA) and neuraminidase (NA) for which the phase affinity has been highly disputed in literature (15, 41–44). Association of viral proteins with raft domains has been difficult to discern, mainly because raft domains are highly dynamic and of submicron size thus making them difficult to be directly visualize in living cells. While HA has been reported to associate with rafts by cell surface analysis (45), it has also been shown to partition into non-raft domains of model membranes (32). However, the lipid preference of HA may be determined by the presence or absence of a palmitoyl anchor which could be responsible for recruiting it to ordered and raft-like phases (46). Here, we used an engineered plasmid to transiently transfect HEK293T cells to express a full-length HA (HA-FL) (Fig. 2) and Fig. S2 which has its C-terminal tagged with a green fluorescent protein (GFP) and is predominantly found to form trimers (47). Also, we engineered a plasmid encoding for HA transmembrane domain with its N-terminal tagged with GFP which is additionally found to form both monomers and dimers (47). A similar strategy was followed for NA (36, 48) and the GPI linked GFP control.

**Fig. 2.**
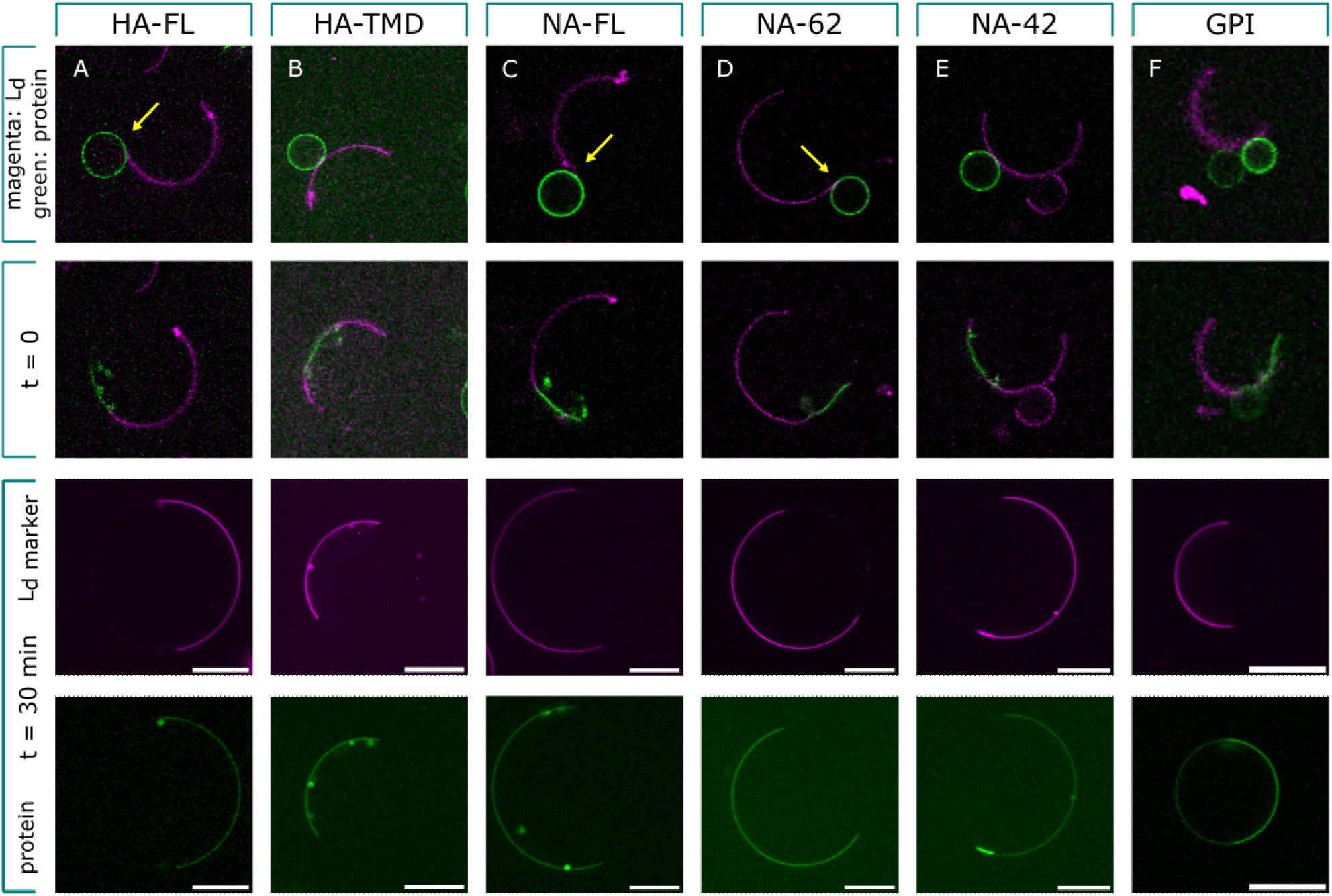
Phase preference of influenza HA, NA and a GPI anchor control. The GPMVs extracted from the plasma membrane accommodate the GFP-tagged protein of interest on their membranes. The disordered phase of the GUVs is marked with the Ld marker 18:1 Liss Rhodamine PE that exclusively partitions into the disordered region. The first row shows the positions where the fusion happens. Yellow arrows indicate where the fusion happens at the ordered-disordered phase interface. At time zero (t = 0), the protein is mainly found near the site of fusion. The phase preference of all proteins was evaluated after 30 minutes (lowest two rows). **A.** Full-length HA protein and **B.** its transmembrane domain co-localize with the liquid disorder phase. **C.** Full-length NA and two C-terminally truncated variants (NA-Δhead) **D.** and (NA-TMD) **E.** distribute into the disordered region. **F.** GPI anchored GFP used as the positive control for liquid order phase partitioning (51). Scale bar is 15 *μ*m.

Similarly to what has been reported before for recombinant HA reconstituted into phase segregated vesicles (49), we find that HA segregates in lipid disordered domains in our hybrid vesicles after fusion. Therefore, contrary to the widely accepted theory of influenza virus assembly (1), HA expressed alone in HEK293T cells does not concentrate in cholesterol enriched lipid domains.

To exclude the possibility that the GFP label attached to the intracellular part of the HA could interfere with the palmitoylation sites we tested the phase preference of the HA transmembrane domain (HA-TMD) with the exctodomain tagged with a GFP. However, as shown in Fig. 2B, the HA-TMD also exhibited complete partitioning into the disordered phase, thus ruling out any interfering effect of the GFP label.

Influenza NA (NA-FL) also displayed a preference for disordered domains and it was primarily driven by the transmembrane domain as similar results were observed using NA constructs lacking the enzymatic head domain (NAΔhead) and the head domain along with the majority of the stalk region (NA1-42), see Fig.2C-E.

As a control we investigated the partitioning of a GPI anchored GFP which has been found to localize to liquid ordered phases in GPMVs (50). As expected GPI predominantly localized into liquid ordered region as shown in Fig.2F.

Quantification of partition coefficients was not possible due to complete partitioning of all proteins apart from the GPI control. As shown in Fig.3 all disordered phases (apart from the GPI experiment) contained no fluorescent signal after subtraction of the image background. Hence, we conclude that, within the sensitivity of our imaging system, both HA and NA exhibit complete partitioning into liquid disordered phases whereas for the GPI control we found a slight Lo preference equivalent to a partition coefficient *K_p,raft_* = 1.4. Definition of *K_p,raft_* can be found in (50).

**Fig. 3.**
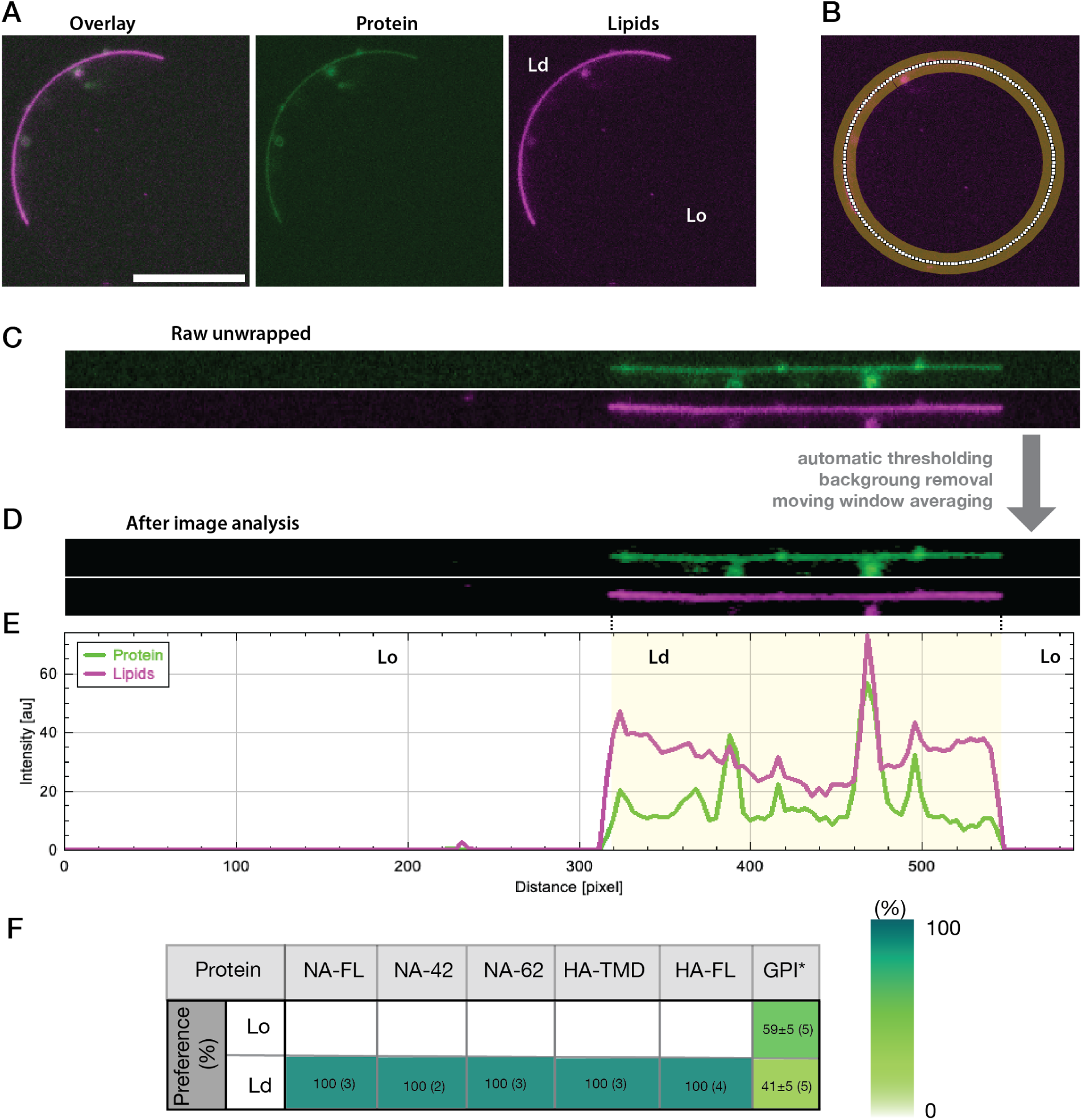
Quantification of protein partitioning. A) Raw data from fusion experiment. Protein (in green) and membrane (magenta) channels are overlayed. Scale bar is 15*μ*m. B) Image exemplifying the selection of the vesicle by the image processing workflow. C) Raw, unwrapped vesicle fluorescent channels. D) Unwrapped vesicles after processing with image processing workflow. E) Fluorescent intensity from the unwrapped vesicles from D). Ld region is highlighted in yellow. F) Lipid order preference from each of the proteins used in this project. We quantified the percentage of protein fluorescence signal coming from Lo and Ld lipid regions. The number in parenthesis indicates the number of fusions performed for each type of protein. * denotes Lo preference.

Our results show that both HA and NA do indeed localize into disordered domains and it remains uncertain whether other effects like protein-protein interactions, between different types of virus proteins, could change this phase preference.

Reports on partitioning of virus proteins into L_o_ or L_d_ phases have yielded conflicting results. Early reports found that the transmembrane domain of HA interacted with sphingolipids and cholesterol as was found by isolating detergent insoluble fractions from the plasma membrane (52), however, the reliability of the detergent resistant method has been questioned (53, 54). Using immuno-electron microscopy together with gold labeling HA was shown to form nanoclusters in fixed cellular plasma membranes. However, no association between HA and rafts was found since it still clustered after depletion of the two raft constituents glycosphingolipids and cholesterol (43). These methods entail procedures that capture raft associated proteins in a single time frame and not in a real-time manner. Other studies have found that HA partitions into raft phases only when a palmitoylation site is present (32). Palmitoylation of proteins has been found to be a major factor for recruiting integral membrane proteins to ordered domains (2, 11, 27, 46). However, this ordered phase recruitment of HA, mediated by the palmitoylation sites, is a mechanism which is not supported by our data. One scenario is that the cytoplasmic GFP label could interfere with the function of the palmitoylation anchor for the FL-HA. However, the HA-TMD which has its GFP label on the ectodomain still partitioned exclusively into the disordered phase. Therefore, we conclude that the palmitoylation site in HA is not sufficient to direct localization of the HA protein into ordered phases.

To avoid any changes to the plasma membrane proteins, GPMVs were derived from cells using N-Ethylmaleimide (NEM) rather than the reducing agent dithiothreitol (DTT); the latter is known to cut thioester-bound fatty acids and therefore, affect the raft-targeting feature of the associated protein. In any case, it remains uncertain how important the palmitoylation is for localizing integral proteins to ordered domains (11) and certainly the hydrophobic match of the transmembrane domains with the thickness of the hydrophobic interior of the membrane is likely to play a role as well (55). Moreover, it should be kept in mind that the cortical actin structure has also been found to organize membrane proteins into clusters (11, 25) and even affect the formation of Lo and Ld phases (42) thus underlining that the membrane organization is not the sole factor determining membrane protein distribution.

More recently (56), a complex interaction was reported with the two influenza matrix proteins during viral budding. These findings, together with our results, suggest that there could be more complex interactions at play during viral budding than previously anticipated (57).

Delivering plasma membrane proteins to GUVs has the advantage that the lipid composition can be controlled and therefore, the hydrophobic mismatch at the Lo-Ld border can be tuned. In addition, since the fused GPMVs are derived from transfected cells they contain the lipid and protein diversity of the plasma membrane although in a diluted version. Importantly the final hybrid vesicle contains several types of proteins correctly oriented as seen from a biological perspective with ectodomains pointing outwards. In addition, the method avoids the need for any protein purification procedures.

Finally, we note that cellular raft domains are dynamic entities allowing both proteins and lipids to move into and out of these domains with different partitioning kinetics thus making rafts difficult to study by direct methods. Although in silico analysis are promising and give a dynamic view of raft kinetics, the results need to be further confirmed by experiments. This highlights the need for a new approach, as presented here, to understand how membrane associated proteins operate in their physiological environment.

## Concluding remarks

Further studies will reveal how coexistence of other proteins from the virus embedded in a model membrane affect their preference for a specific lipid phase. An extremely interesting future perspective of the current method is the possibility of combinatorial selection of proteins to be added sequentially to a hybrid GUV/GPMV model membrane. Influenza virus proteins HA, NA, M1 and M2 proteins are populating the virus envelope and our assay can be employed to systematically investigate possible inter-protein interactions which could change their phase affinity and even other membrane remodeling behavior that these proteins could be responsible for. In general our method allows for generic investigation of how single proteins, and mixtures of proteins, collectively organize in heterogeneous membranes which will advance our understanding of protein function in complex biological membranes.

## ACKNOWLEDGEMENTS

This work is financially supported by Danish Council for Independent Research, Natural Sciences (DFF-4181-00196) and Novo Nordisk Foundation Interdisciplinary Synergy Program 2018 (NNF18OC0034936). We wish to thank Henrik Østbye who was involved in the development of the NA and NA-mutant plasmids as previously reported in (36). We will also like to acknowledge Nicola De Franceschi for fruitful discussions about this project back at Patricia’s Bassereau lab at the Curie Institute.

## Materials and Methods

### Cell Culture

Human Embryonic Kidney Cell line (HEK293T) was cultured in Dulbecco’s Modified Eagle Medium (DMEM, Gibco, cat. no. 11995065) supplemented with 10% fetal bovine serum (FBS, Gibco, cat. no. 11550356) and 100 U/mL penicillin and 100 mg/mL streptomycin (Gibco, cat. no. 15140122). The cells were grown at 37 °C in 5% CO_2_ in a humidified incubator.

### Cell Transfection

3 × 10^5^ HEK293T cells were seeded on a 35mm glass bottom dish (MatTek, cat. no. P35G-1.5-14-C) coated with 0.01% poly-L-lysine (Sigma-Aldrich, cat. no. P8920) and grown for 24 h. Then, the cells transiently transfected with the plasmid of interest using Lipofectamine™ LTX Reagent (Invitrogen, cat. no. 15338030) according to the manufacturer’s protocol with minor optimization to improve expression yield.

### Giant Plasma Membrane Vesicle (GPMV) Preparation

GPMVs were obtained according to (1). In brief, transfected cells were grown for 24 h and then washed with PBS (Gibco, cat. no. 10010023) and GPMV buffer (10 mM HEPES, 150 mM NaCl, 2 mM CaCl2, pH7.4) in succession. The vesiculation process triggered by adding to the cells 1 mL GPMV buffer to which N-Ethylmaleimide (NEM, Sigma-Aldrich, cat. no. E3876) was added to the final concentration of 2 mM. After 2 h of incubation at 37 °C, GPMVs were ready for the experiment.

### Giant Unilamellar Vesicle (GUV) Preparation

GUVs were made according to the PVA gel-assisted hydration method (2) using a mixture of DOPC (1,2-dioleoyl-sn-glycero-3-phosphocholine, Avanti Polar Lipids, cat. no. 850375), Brain SM (Avanti Polar Lipids, cat. no. 860062) and Cholesterol (Avanti Polar Lipids, cat. no. 700000) dissolved in chloroform with a molar ratio of 2:2:1. Furthermore, the lipids were mixed with the Ld marker 18:1 Liss Rhod PE (1,2-dioleoyl-sn-glycero-3-phosphoethanolamine-N-lissamine rhodamine B sulfonyl, Avanti Polar Lipids, cat. no. 810150) at 0.5 mol.%. For the GUV-GPMV fusion experiments, 1 mol.% of DSPE-PEG(2000) Biotin (1,2-distearoyl-sn-glycero-3-phosphoethanolamine-N- [biotinyl(polyethylene glycol)-2000], Avanti Polar Lipids, cat. no. 880129) were also added to the lipid mix.

### Optical Tweezers and Fusion

A Leica SP5 confocal microscope implemented with an optical trap based on a 1064 nm laser (Spectra Physics J201-BL-106C)) were used for confocal visualization and optical trapping (3). A Leica PL APO 63X water immersion objective with NA = 1.2 was used for sample visualization and focusing the trapping laser beam. The GFP tag and Rhodamine in the sample were excited with 488 nm and 594 nm laser lines respectively and their emitted intensities were collected in the spectral range of 493-553 nm and 598-700 nm. The trapping laser operated with the output power of 45 mW corresponding to an intensity of 3 × 10^10^ W m^-2^ in the sample and irradiation of Streptavidin-coated 150 nm AuNSs (nanoComposix, cat. no. GSIR150) in the interface of a GUV-GPMV triggers the fusion event. Also, to spot the AuNSs a 476 nm argon laser line was used and the reflected intensity was captured in the spectral range of 476-488 nm.

### Design and Preparation of Plasmids

Plasmid design and preparation was described in (4). *pCAG*: *GPI — GFP* was a gift from Anna-Katerina Hadjantonakis (Addgene plasmid #32601, http://n2t.net/addgene:32601., *RRID*: *Addgene_32601_*)

### Data Analysis

Image processing was all carried out in Fiji (5, 6), using macro language commands for unwrapping the vesicles signals and extracting the fluorescence intensities along the membrane. Fluorescence intensity curves where then plotted and analyzed using Python. Macro image workflow can be found in the following Github repository https://github.com/GMorenoPescador/vesicle_unwrapper.

## Supplementary Information

**Fig. S1.**
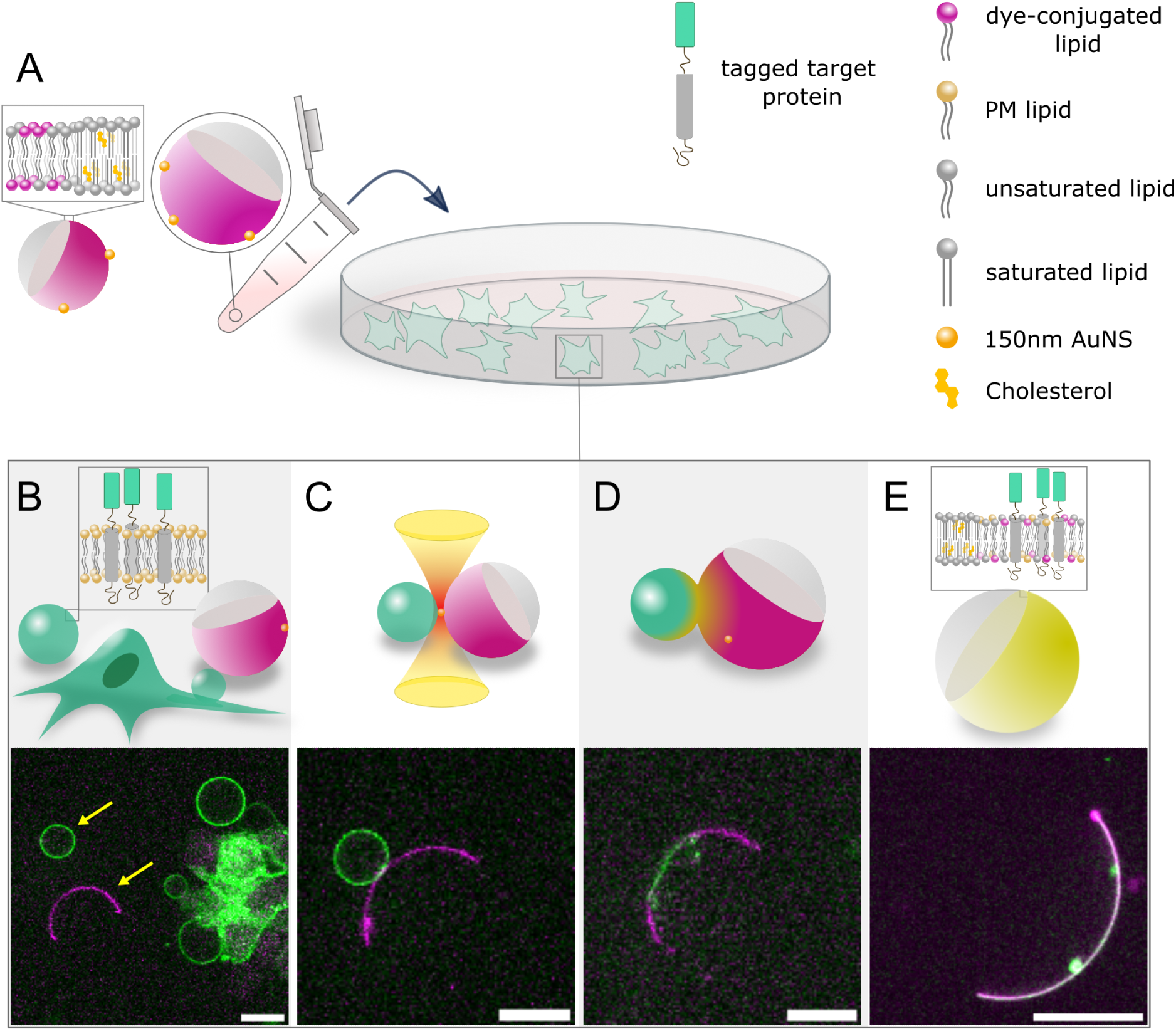
Systematic delivery of membrane associated proteins into phase-separated GUVs. **A.** HEK293T cells are transfected to express the protein of interest that itself is a chimeric fluorescent protein, here GFP-tagged Hemagglutinin Transmembrane Domain (GFP-HA-TMD). The cells then treated with a vesiculating agent which causes cells to exclude part of their plasma membranes as GPMVs that accommodate the expressed protein on their membranes. Phase-separated GUVs are also added to the culture dish. **B.** As a result, the culture dish gets populated with GPMVs and GUVs with varying sizes that enables target selection from a pool of vesicles. A GPMV is then grabbed by the optical trap and brought into close contact with a GUV (yellow arrows). **C.** Irradiating the interface of the GPMV-GUV pair with the NIR laser pulls GUV-bound gold nanoshells into the laser focus where they produce highly localized heat. **D.** The produced heat melts the opposing membranes and fusion happens. **E.** The resultant hybrid vesicle inherits its lipid composition from the GUV-GPMV and retains its phases separated. In this way, the protein of interest has been transferred from the cell plasma membrane to the model membrane where it redistributes in its preferred phase - that is disordered phase in this example. Insets in A, B and E show the membrane composition of each vesicle. Scale bars are 15 μm.

**Fig. S2.**
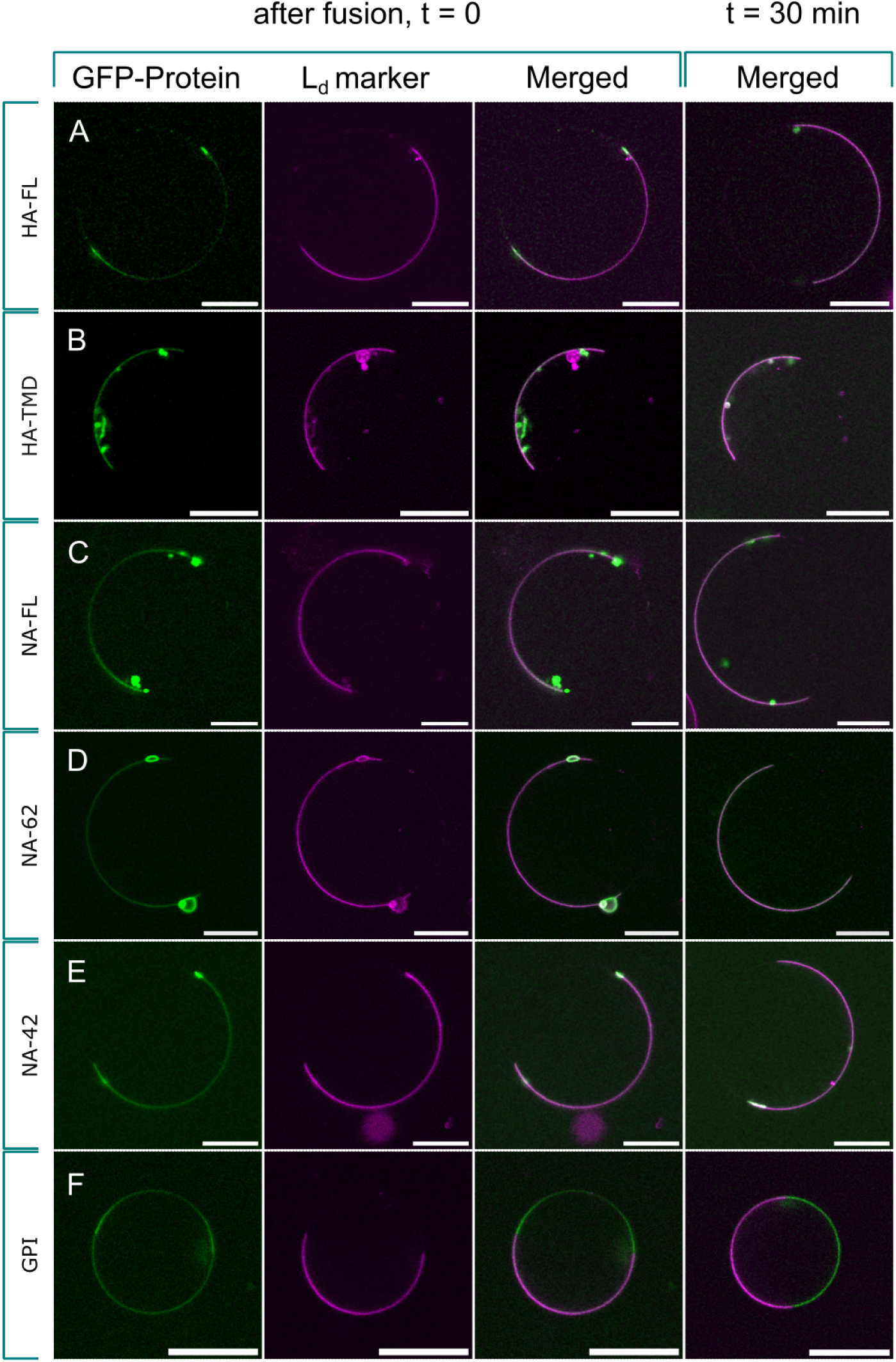
Phase partitioning of influenza A virus transmembrane proteins. When the protein of interest is delivered into the phase-separated GUV, it starts to distribute on it. After 30 minutes of the fusion event, the hybrid vesicle still retains its phases separated and the protein is found on its preferred phase. The disordered phase is labeled with 18:1 liss. rhodamine PE lipid that exclusively partitions into the liquid-disordered region (colored in magenta). **A.** Full-length Hemagglutinin (HA-FL). **B.** Transmembrane domain of Hemagglutinin (HA-TMD). **C.** Full-length Neuraminidase (NA-FL). **D.** Truncated Neuraminidase that lacks the bulky head domain (NA-62). **E.** Another truncated Neuraminidase that lacks the bulky head domain and also has a shorter stem region (NA-42). NA-42 forms dimer instead of tetramer on the membrane. **F.** GPI anchored GFP as the positive control of true partitioning. Scale bars are 15 μm.

