## Supplemental text and figures for "Thermoplasmonic induced vesicle fusion for investigating membrane protein phase affinity"

### Supplementary Information

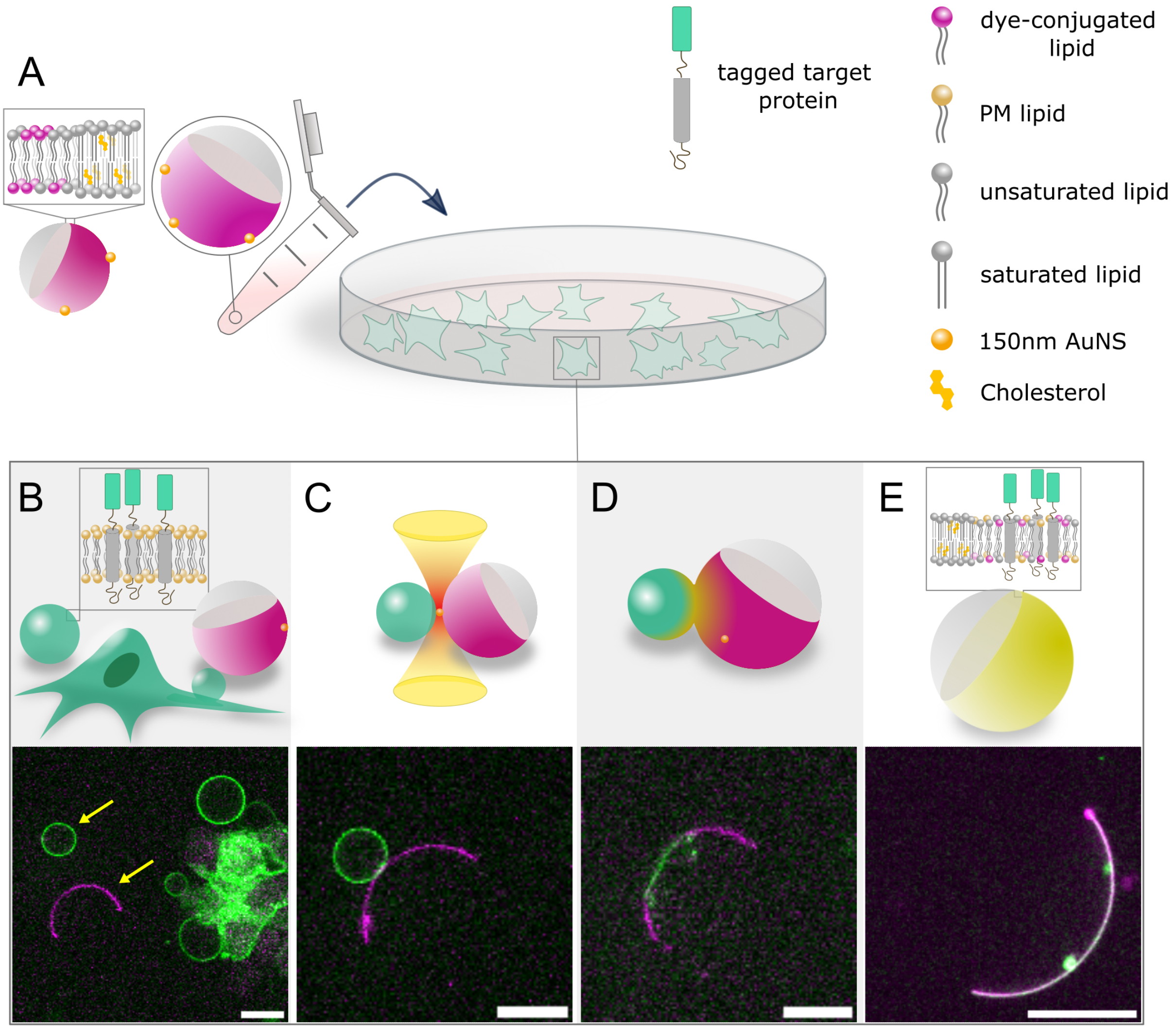


**Fig. S1. Systematic delivery of membrane associated proteins into phase-separated GUVs. A.** HEK293T cells are transfected to express the protein of interest that itself is a chimeric fluorescent protein, here GFP-tagged Hemagglutinin Transmembrane Domain (GFP-HA-TMD). The cells then treated with a vesiculating agent which causes cells to exclude part of their plasma membranes as GPMVs that accommodate the expressed protein on their membranes. Phase-separated GUVs are also added to the culture dish. **B.** As a result, the culture dish gets populated with GPMVs and GUVs with varying sizes that enables target selection from a pool of vesicles. A GPMV is then grabbed by the optical trap and brought into close contact with a GUV (yellow arrows). **C.** Irradiating the interface of the GPMV-GUV pair with the NIR laser pulls GUV-bound gold nanoshells into the laser focus where they produce highly localized heat. **D.** The produced heat melts the opposing membranes and fusion happens. **E.** The resultant hybrid vesicle inherits its lipid composition from the GUV-GPMV and retains its phases separated. In this way, the protein of interest has been transferred from the cell plasma membrane to the model membrane where it redistributes in its preferred phase - that is disordered phase in this example. Insets in A, B and E show the membrane composition of each vesicle. Scale bars are 15 µm.

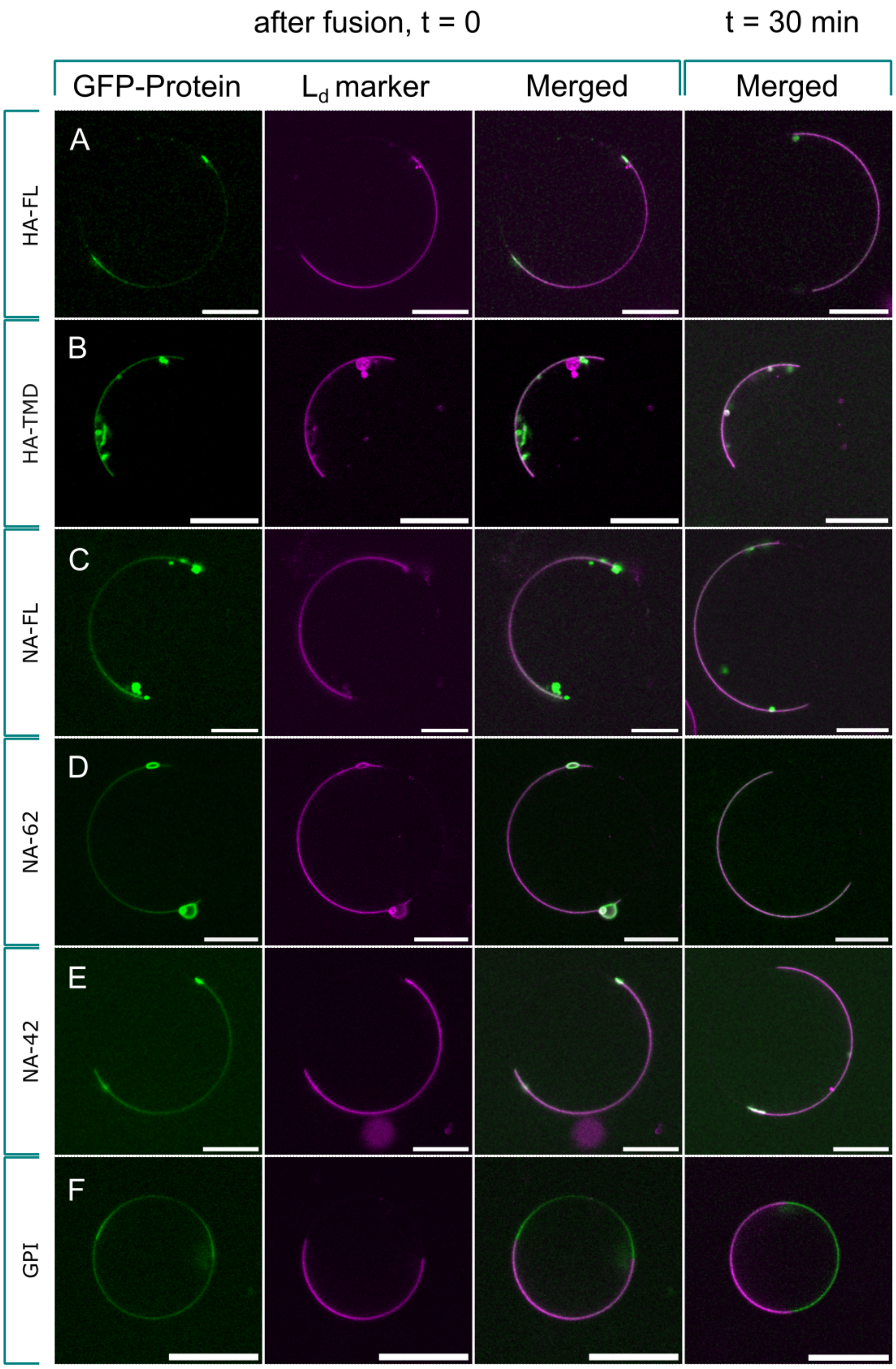


**Fig. S2. Phase partitioning of influenza A virus transmembrane proteins.** When the protein of interest is delivered into the phase-separated GUV, it starts to distribute on it. After 30 minutes of the fusion event, the hybrid vesicle still retains its phases separated and the protein is found on its preferred phase. The disordered phase is labeled with 18:1 liss. rhodamine PE lipid that exclusively partitions into the liquid-disordered region (colored in magenta). **A.** Full-length Hemagglutinin (HA-FL). **B.** Transmembrane domain of Hemagglutinin (HA-TMD). **C.** Full-length Neuraminidase (NA-FL). **D.** Truncated Neuraminidase that lacks the bulky head domain (NA-62). **E.** Another truncated Neuraminidase that lacks the bulky head domain and also has a shorter stem region (NA-42). NA-42 forms dimer instead of tetramer on the membrane. **F.** GPI anchored GFP as the positive control of true partitioning. Scale bars are 15 µm.
